# Occurrence and quantities of DNA modifications across the tree of life

**DOI:** 10.1101/2022.03.22.485282

**Authors:** Sreejith Jayasree Varma, Enrica Calvani, Nana-Maria Grüning, Christoph Messner, Nicholas Grayson, Floriana Capuano, Michael Mülleder, Markus Ralser

**Author notes:** To whom correspondence should be addressed. M. Ralser, Charité Universitätsmedizin Berlin. These authors contributed equally to the work (Sreejith Jayasree Varma, Enrica Calvani).

## Abstract

Enzymatic DNA modifications like methylcytosine (5mdC), methyladenine (N6mdA), or hydroxymethylcytosine (5hmdC) are key for chromatin function, gene expression regulation, and antiviral defense, but they remain understudied in non-model organisms. We established a mass spectrometric method for the sensitive and accurate quantification of enzymatic DNA modifications, and analyzed 85 bacterial genomes, 19 plant samples, 41 tissues from 12 animal species, 6 yeast species, and two archaeal species. We report no or only very low concentrations of DNA modifications in yeast and insects, but find DNA modifications universal to both bacteria and higher eukaryotes. Specifically for prokaryotes, our dataset indicates that evolutionary relationships and host–pathogen interactions, but not the ecological niche in general, select for a similar degree of DNA modification. In higher eukaryotes, largest concentration differences between tissues are detected for 5hmdC. Our dataset further reveals unique biological cases that warrant attention in the study of DNA modifications. For instance, while our data shows that most species contain just one dominating DNA modification, we detect all dominianting DNA modifications (5mdC, N6mdA, and 5hmdC) to exist in parallel in *Raphanus sativus*. Other plant species, like onion, sunflower, or the grass big bluestem, can have more than 35% of cytosines methylated. Finally, 5hmdC, so far mostly studied in the vertebrate central nervous system, was identified to reach a concentration of up to 8% of all cytosines in the Oman garra brain, and was also detected in several plants, like *Lepidium sativum*. The present study underscores the exploitation of biological diversity for studying DNA modifications.

## Introduction

Enzyme-catalyzed DNA modifications are studied for their roles in chromatin structure, gene-expression regulation, prevention of viral DNA integration, epigenetic inheritance, cell– environment interactions, developmental biology, immunity, memory, aging, and cancer (1–10). The methylation of the 5th carbon (C5) of the cytosine ring to yield 5-methyl-2′-deoxycytidine (5mdC) was the first nucleotide modification to be discovered (11), and has remained the most intensively studied (12,13). 5mdC can be enzymatically oxidized into 5-hydroxymethyl-2′-deoxycytidine (5hmdC) and further into 5-formyl-2′-deoxycytidine (fdC) and 5-carboxyl-2′-deoxycytidine (cadC) (14–16). Although these modifications have been described as transient intermediates of 5mdC demethylation, at least one (5hmdC) has been found to accumulate in the mammalian brain, specifically in the large Purkinje neurons, indicating a regulatory function (17). N4-methyl-2′-deoxycytidine (4mdC), found in bacteria, is yet another form of cytosine modification (18,19). Cytosine thus exists in multiple chemical states (dC, 5mdC, 5hmdC, fdC, cadC, 4mdC, as well as the rare 4,5-dimethyl-2′-deoxycytidine (4,5dmdC)) (12,20). Another important modification is the N6 methylation of adenine. N6-methyl-2′-deoxyadenosine (N6mdA) was initially discovered in bacterial genomes (21) and later also in archaea, plants, and nematodes (22,23). Although N6mdA is not essential in microbial model organisms, this modification has been increasingly associated with functions that promote virulence or viral DNA integration (24,25). Indeed, it seems likely that DNA modifications play different roles in different species, as indicated by the varying amounts of DNA modifications across model organisms. For instance, *Arabidopsis thaliana* has orders of magnitude higher levels 5mdC compared to the dominant insect model *Drosophila melanogaster*, while the dominant yeast model organism *Saccharomyces cerevisiae* lacks this modification altogether (26,27).

Since until recently studying DNA modifications was technically challenging, information concerning their content and function is still scarce for species other than model organisms, several crops, and humans. Therefore, it is rather difficult to translate the knowledge derived from those intensively studied species into a broader biological context. For instance, it is hard to judge from the current literature if the low amount of DNA modifications in laboratory yeast and *D. melanogaster*, or the high amount in *A. thaliana (27)*, represent the rule or the exception in their respective phylogenetic group without a broader multi-species dataset for comparison.

Sequencing technologies are the dominating method for studying the role of DNA modifications in humans and model organisms. They provide only relative quantitative values about the consent of DNA modifications, but position-specific information, which is required to understand the specific molecular functions of modification, for instance whether they influence gene-expression regulation in a specific locus (28,29). On the other hand, it is equally important to absolutely quantify DNA modifications at the genome-wide level. Quantitative values help to address more-general questions, like evolution and activity of the biochemical pathways that modify nucleic acids, their role in viral immunity, relationship between different modifications, and for comparing (non-model) species still lacking high-quality reference genomes. We and others (27,30–34) have shown previously that targeted mass spectrometry is an ideal technology to determine absolute quantities of DNA modifications, specifically, if they are low abundant and in the noise range of sequencing technologies. Mass spectrometry further is suitable for studying poorly characterized species, as no prior knowledge about the genome is required for data analysis. Aside from that, targeted mass spectrometry is economical, with running costs per sample amounting to single-digit dollars. For these reasons, mass-spectrometric quantification is well suited for identifying interesting patterns in the amount and relative abundances of DNA modifications, specifically within understudied species.

## Results and discussion

### Quantification of a panel of enzymatic DNA modification using liquid chromatography/multiple reaction monitoring

In order to quantify multiple enzymatic DNA modifications in a single analysis, we expanded a previous method based on liquid chromatography–multiple reaction monitoring (LC–MRM) and designed for the quantification of 5mdC (35). This method is characterized by a sensitivity down to attomoles and a broad dynamic range, and discriminates between RNA and DNA modifications, clarifying the previously debated content of 5mdC in several yeast species (27). In this method, isolated DNA is first enzymatically digested to obtain the corresponding nucleosides using a nuclease enzyme mixture (DNA Degradase Plus, Zymo Research). The resulting digests are then filtered and directly analyzed by a targeted assay using liquid chromatography - multiple reaction monitoring (LC–MRM) using a triple quadrupole (QQQ) mass spectrometer. Distinguishing the nucleosides arising from a DNA monomer from a potentially co-purified RNA monomer occurs on the basis of the precursor mass difference of the sugar moiety. Such a strategy ensures the measured nucleosides are free from RNA contamination as many base modifications are more frequently observed in RNA (27,35). For quantifying other DNA modifications, namely 5hmdC, N6mdA, cadC and fdC, we obtained synthetic standards for these molecules (Methods, Table S1) and optimized the instrumental and chromatography parameters accordingly. Moreover, we supplemented the method by a neutral loss scan to confirm the MRM results, as well as to detect additional modifications like 4mdC, or to screen for yet unknown modifications (Methods). Combined with the high sensitivity offered by a triple quadrupole mass spectrometer (Agilent 6470), we were able to achieve detection limits in picomolar ranges (Fig. 1A).

**Figure 1.**
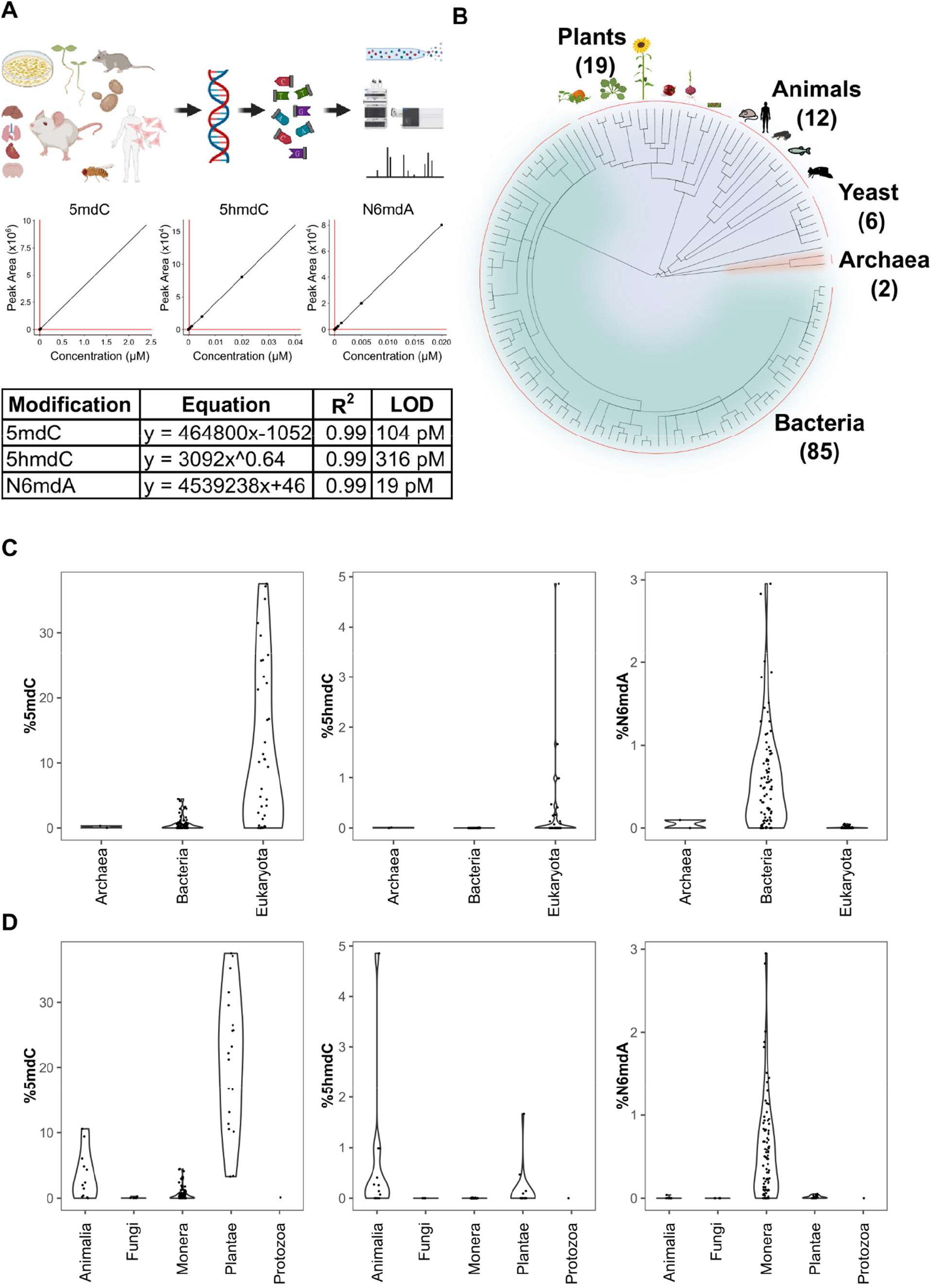
(A) Multiplex analysis of various genomic DNA modifications using liquid chromatography multiple reaction monitoring (LC-MRM) following enzymatic digestion of DNA. The regression curves and limit of detection (LOD) for modifications 5mdC, 5hmdC, and N6mdA are represented. Although our method also quantifies cadC and fdC, we did not detect significant concentrations of these in any of the measured samples; these modifications were hence omitted from the graphical illustrations. (B) 286 samples from 124 species were analyzed in the present study: 19 species from plants, 12 from animals, 6 from yeast, 2 from archaea, and 85 from bacteria. (C–D) Distribution of 5mdC, 5hmdC, and N6mdA across (C) archaeal, bacterial, and eukaryotic domains, and (D) animal, fungi, monera, plant, and protozoan kingdoms. The values depict percentage of cytosine residues bearing either methyl (%5mdC) or hydroxymethyl (%5hmdC) modification and percentage of adenine residues bearing methyl modification (N6mdA).

Upon setting up the method, we sampled cells or tissues for a large number of species across the three domains of life. Because our method does not include any amplification steps, and detects modifications on the DNA directly, it requires clean DNA at microgram levels, at least for the detection of the lowly concentrated DNA modifications. Unfortunately for some rare specimens, we only had limited sample amounts, and in many cases, standard DNA preparation protocols did not yield DNA of sufficient quality or concentration for our assay. However, by combining different protocols, we were able to extract clean DNA at microgram levels for 286 distinct cases. These are derived from 124 different species, including 85 bacterial species, 6 yeast species, 2 archeal species, 19 plant species, and 18 tissue and cell-culture samples from 12 animal species, including human and mouse. The collection included both the typical model organisms, and specifically for bacteria, vertebrates, and plants we included a significant number of species that have been barely characterized at the molecular level so far (Fig 1B). Furthermore, for a number of vertebrates, including human, the model organisms mouse (*Mus musculus)*, African clawed frog (*Xenopus laevis*), but also for some less studied species, the opossum *(Monodelphis domestica*), the Alpine marmot (*Marmota marmota*), and the Oman garra (*Garra barreimiae*), we obtained DNA from multiple tissues and/or cell lines in order to quantify tissue differences in the absolute DNA modification content. For plants, we focused on seedlings that were germinated in the lab (Methods). The seedlings not only allowed for efficient DNA extraction, which can be hampered by high concentrations of plant polymers in fully differentiated plant tissues, but also for direct comparison between the plants at a similar developmental stage.

### While multiple lower eukaryotes lack DNA modifications, N6mdA dominates in bacteria, and 5mdC is the dominating DNA modification across higher eukaryotes

Our results reveal major global differences in the nature and total concentration of DNA modifications when comparing the domains of life (Fig. 1C–D). First, despite the broad coverage, high sensitivity, and precision of our method, we did not detect significant levels of fdC and cadC in any of the genomes measured (limits of detection were 238 pM and 251 pM, respectively). These oxidized forms of 5-methyl-2′-deoxycytidine have been associated with the degradation of 5mdC (15), and according to our results they seem to remain transient across species as they do not accumulate to significant, genome-wide-scale levels. Next, we detected hardly any DNA modification in any of the unicellular fungi studied (Table S3). Hence it is not merely 5mdC (27,36,37), but also its oxidized form 5hmdC along with N6mdA that are very low if not absent in typical yeast species. Is interesting in this context that the insects *Trichoplusia ni, Spodoptera frugiperda and D. melanogaster* (Table S3) all had very low amounts of these DNA modifications as well. Indeed, the fruit fly *D. melanogaster* has so far been considered an unusual case among the laboratory model organisms, as it contains only trace amounts, if any, of cytosine methylation (27,38,39). The presence of other DNA modifications in *D. melanogaster* like N6mdA has also been contested due to the presence of an appreciable gut microbiome, which could confound the results (40). We assessed this situation, comparing the genomic DNA obtained from fruit flies that possessed a functioning gut microbiome versus ones grown under germ-free conditions. N6mdA was also detected in germ-free *D. melanogaster* (∼0.04%, Fig. S1). In a recent study comparing DNA adenine methylation levels in multiple eukaryotic species, the bacterial contamination affected the N6mdA measurements. However, it was possible to distinguish the N6mdA in *Drosophila* tissue from microbial contamination using quantitative deconvolution (41). While the adult *D. melanogaster* contained methylated adenine as a DNA building block, ovarian cells collected from two moth species (*T. ni* and *S. frugiperda*) principally contained methylated cytosine as the preferred base modification (0.2% and 0.1%, respectively).

Which conclusions can be drawn from the low concentrations of DNA modifications in yeasts and insects? First, these results support the notion that enzymatic DNA modifications are not universal, which could have peculiar evolutionary consequences. Studies in yeast have concluded that DNA modifications could have been specifically lost during yeast evolution (42). However, our result that insects can have similarly low DNA modification levels raises another possibility: that DNA modifications could have evolved in higher eukaryotes and bacteria, after yeasts and insects branching off.

As a rule, most genomes contained a single modification type. Some exceptions to this were however encountered. A subset of the eukaryotes and a subset of prokaryotic species contained low concentrations also of a second modification, which could be either 5mdC, N6mdA, or 5hmdC (Fig. 2, Table S3, Fig S2). For instance, *Diplotaxis tenuifolia* had low amounts of N6mdA (0.1%, Table S3) next to high amounts of 5mdC. Of particular interest was *Raphanus sativus*, which was the only species among those analyzed that possessed all the three modifications at detectable levels and in parallel. Among prokaryotes, we observed a maximum of two modifications (5mdC and N6mdA), with 5hmdC entirely missing. Our study further featured two archeal genomes (*Sulfolobus acidocaldarius* and *Halobacterium salinarum*), which shared a similar level of the cytosine modification but differed in their levels of adenosine modification. While we detected N6mdA in *Halobacterium*, no adenosine modification was observed for *Sulfolobus* (Table S3).

**Figure 2.**
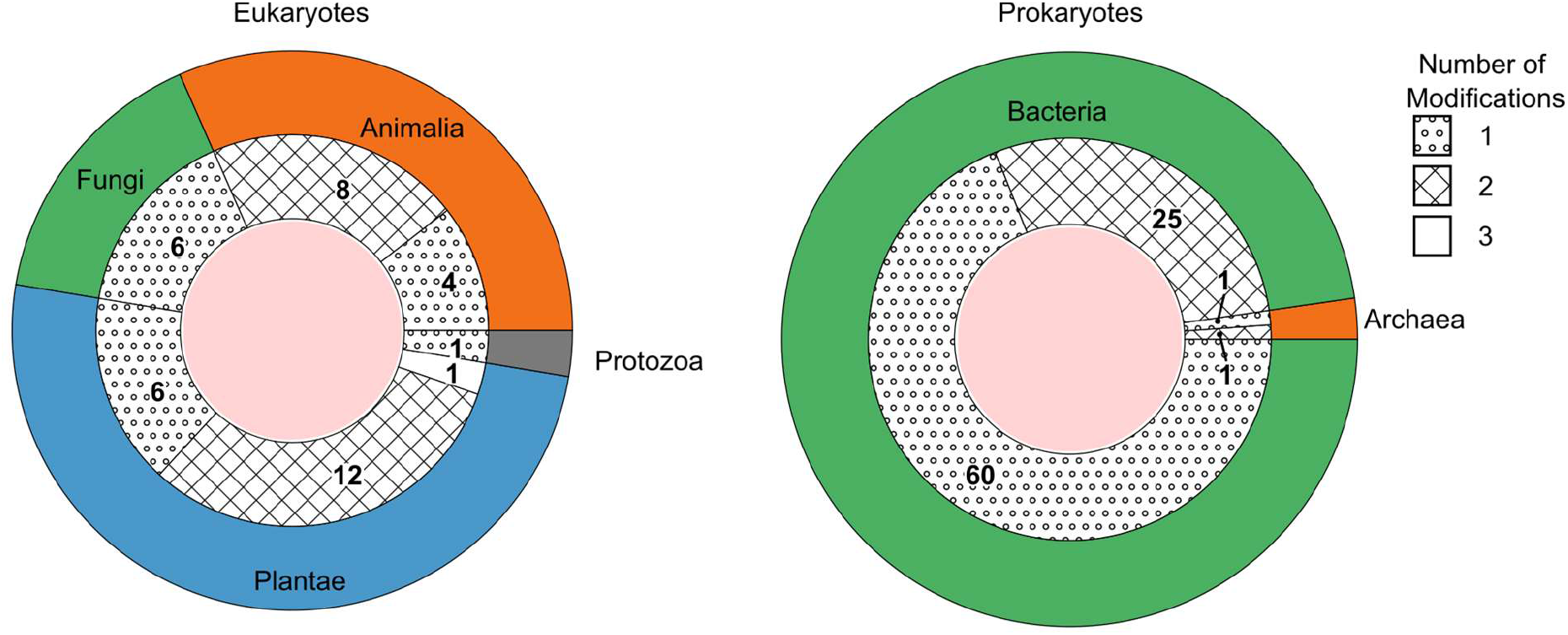
The number of species possessing one, two, or three DNA modification types grouped as eukaryotes (left) and prokaryotes (right). The outer ring represents the kingdoms present within these domains. The groupings per number of modifications are shown as fill patterns on the inner ring, where dots represent species in which only one among 5mdC, 5hmdC, and N6mdA were found; crosses represent species bearing two modifications simultaneously; and no fill represents species carrying all three modifications.

### Tissue divergence of 5mdC concentrations in vertebrate and plant genomes

Among the DNA modifications, 5mdC had the highest abundance and was specifically abundant in plants. Most vertebrate genomes studied had a 5mdC content of around 5% (mean 4.66, SD 2.17) of the cytosine residues. Some species, including the model organisms *Danio rerio* and *Xenopus laevis*, had higher levels consistent with early observations (43). In plants however, 5mdC concentrations of 10% (mean 20.34, SD 9.81) and higher were typical (Fig. 1D). Extremely high values for cytosine methylation were observed in *Andropogon gerardii* and *Allium cepa*, where more than 35% of cytosines were methylated (Fig. 1D, Table S3). Given that very low levels or no 5mdC were detected in yeast and insects, cytosine 5 methylation levels hence differ by several orders of magnitude within the eukaryotic kingdom.

In multicellular organisms, DNA modifications are important for development, and tissue differences between DNA modification patterns are observed (44–46). Our data suggests that a change in the modification pattern or sequence context does not necessarily have a strong impact on the total concentrations of the DNA modifications however. We analyzed spleen, muscle, lung, liver, kidney, heart, and central nervous system (CNS) samples from five animal species, of which two are model organisms *(Xenopus laevis, Mus musculus)*, and three non-model organisms *(Garra barreimiae, Monodelphis domestica, Marmota marmota*). From *M. musculus* we further examined tissues from multiple inbred laboratory lines: BALB/c, FVB/N, Hsd/Ola/MF1, B6SJL/CD451/CD452, BALB/cAnN, 129S8, and F1/CBAxB6. In parallel, we analyzed multiple human cell lines (Table S3). The obtained data was consistent, in the sense that the values for 5mdC levels were highly similar, as long as the tissues were derived from the same species (Fig. 3A, left). For instance, most tissues in *G. barreimiae, M. marmota*, and *M. musculus* tissues had 5mdC levels of around 5–6% (Fig 3A). Between the different mouse lines, there were no significant differences in 5mdC levels (Table S3). We noted, however, some small but notable differences between specific tissues. Heart tissue presented a broad cytosine methylation level and brain tissue had a higher median value for percentage methylation compared to other tissues (5.3% vs 4.9%) (Fig 3B). We then tested whether different nutritional conditions would change the picture. Therefore, we grew a commonly used mammalian cell line (HeLa) under different growth conditions. The different growth conditions affected 5mdC levels, and the detected differences were in a similar magnitude as the small differences detected between tissues (Fig S3).

**Figure 3.**
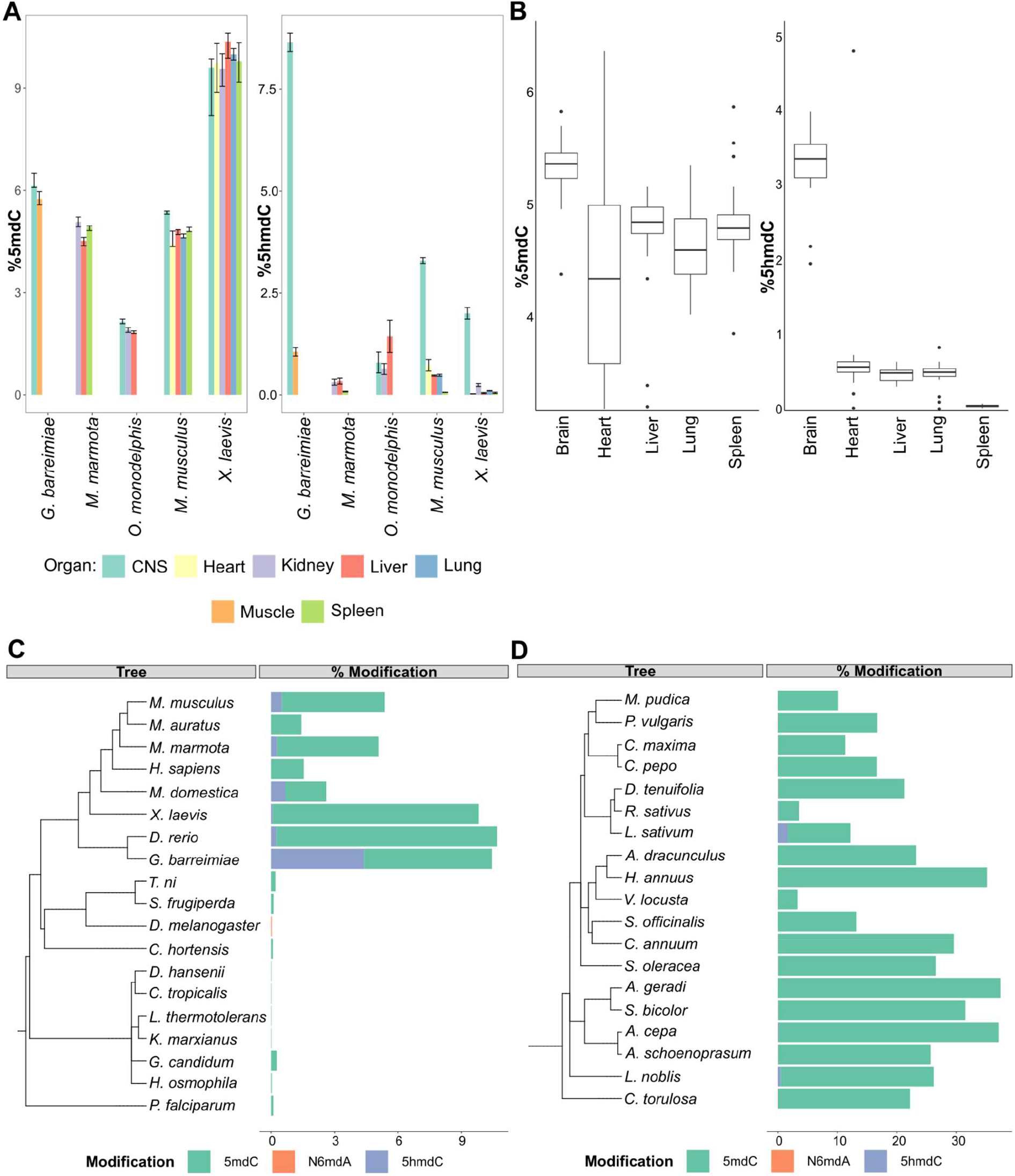
(A) The concentration of 5-methyl deoxycytidine (left) and 5-hydroxymethyl deoxycytidine (right) in different vertebrate genomes. (B) Distribution of 5-methyl deoxycytidine (left) and 5-hydroxymethyl deoxycytidine (right) in different mouse tissues. Variations in percentage modification across different (C) non-plant eukaryotes including representatives from vertebrates like mammals, amphibians, and fish, invertebrates like insects and molluscs, and unicellular fungi and protozoa (D) plants species comprising both gymnosperms and angiosperms.

Overall 5mdC concentrations in opossum and *Xenopus*, respectively, were different to the aforementioned species. In opossum, we detected much lower levels (2%) of 5mdC in all tissues examined. Conversely, in *X. laevis*, all tissues had much higher concentrations (about 9.4%). However, also here, in both cases the tissue differences in the 5mdC concentrations were minimal, at least when compared to the differences that exist between species. Although we tested fewer cases in plants, our data suggest the situation could be similar there too. We tested different tissues (roots, leaf, stem, and seed cotyledon) from *Phaseolus vulgaris*, and obtained consistently high (16.7%) 5mdC concentrations in all measured tissues (Table S3). Hence, the several tissues examined from animal species, cell lines, and *Phaseolus vulgaris* provided a largely consistent picture: in a given organism, several tissues exhibit similar levels of 5mdC, and, that within-tissue differences are typically smaller compared to the differences that can be detected between species.

### Tissue specificity of 5-hydroxymethyl deoxycytidine in the vertebrate CNS

Tissue specificity was, however, detected for another modification, 5-hydroxymethyl deoxycytidine (5hmdC). Indeed, 5hmdC was previously discovered in mammalian brain tissue, where it is formed via oxidation of 5mdC by TET enzymes (16,47). Our dataset shows that 5hmdC is detected in a broad range of vertebrate tissues except for spleen, but reaches significantly higher concentrations specifically in samples from the CNS. Although the spleen tissues had similar 5mdC levels as other mouse tissues, 5hmdC was not detected in these tissues (Fig. 3B). Interestingly, our data reveals that the highest 5hmdC levels were not detected in the mammalian brain. The presence of this modification could reach up to 8% of cytosine residues in *G. barreimiae. M. musculus* (3.3%) and *X. laevis* (2%) too had high levels of 5hmdC specifically in brain tissue relative to other tissues in those organisms (Fig. 3A, right). An interesting exception was in opossum, the only vertebrate species analyzed, in which 5hmdC levels were not higher in the brain compared to peripheral tissue.

Apart from vertebrates, 5hmdC was also observed in *A. thaliana* and *Oryza sativa* (48). Our data shows that the presence of 5hmdC is by no means universal in plants. Indeed we did not detect it in the majority of plant samples. However, our data adds several species (*Allium cepa, Laurus nobilis, Lepidium sativum*, and *R. sativus*) in which we confirmed low concentrations of 5hmdC. Further, we did not detect 5hmdC in any of the bacterial or fungal genomes analyzed. Our results support the fact that the modification of 5hmdC is more widespread in biological systems as previously assumed, but are not universal or specific, to any part of the phylogenetic tree.

### Variations in DNA modification across different bacterial species

In prokaryotes, high amounts of DNA modifications all concerned N6mdA, with the highest levels detected in *Mobiluncus curtisii* (∼1.4%) and *M. thermoacetica* (∼1.1%). In total, the prokaryotic genomes hence contained higher amounts of DNA modifications compared to lower eukaryotes such as yeasts and insects, but lower amounts of DNA modifications compared to higher eukaryotes—plants and vertebrates in particular.

Typical bacterial species contain only one modification type—mostly N6mdA (Fig. 4A). Our data reveals some exceptions. Certain genera such as *Campylobacter* contain trace quantities of 5mdC (<0.1%) next to the dominating N6mdA modification (Table S4). In general, the observed trend was that the occurrence of one type of modification limits the occurrence of the other. For instance, *M. curtisii* with ∼1.4% of its adenine residues methylated shows only 0.3% 5mdC, while *Sebaldella termitidis*, with unusually high cytidine methylation (∼2.4%), has only 0.1% of its adenines methylated. Interestingly, we observed that median values for 5mdC dominate over N6mdA in those bacteria that colonize or enter mutualistic relationships with higher eukaryote species that carry 5mdC as their main modification (Fig. S4, Table S4). This included the genus *Neisseria*, mucosal-surface-colonizing bacteria, which showed 1.4% and 2% (*N. gonorrhoeae, N. lactamica* respectively) of cytosine residues were methylated while containing only <0.3% N6mdA, and *Faecalicoccus pleomorphus* and *Bifidobacterium adolescentis*, with >1.5% of 5mdC without any detectable levels of N6mdA modification. Indeed, others made a similar observation in single-cell fungi. While the environmental yeasts studied lacked any modifications (27), the most frequent commensal yeast pathogen *Candida albicans* contains 5mdC (49). This result is interesting, because it is believed that the main role of DNA modifications in single cellular organisms is a defense against viruses, while higher organisms adapted methylation for gene regulation. In that light, the increase of DNA methylation in the pathogen in host–pathogen interactions would be hard to explain. Our data suggests hence that the picture about the functions of DNA modifications in prokaryotes is not complete.

**Figure 4.**
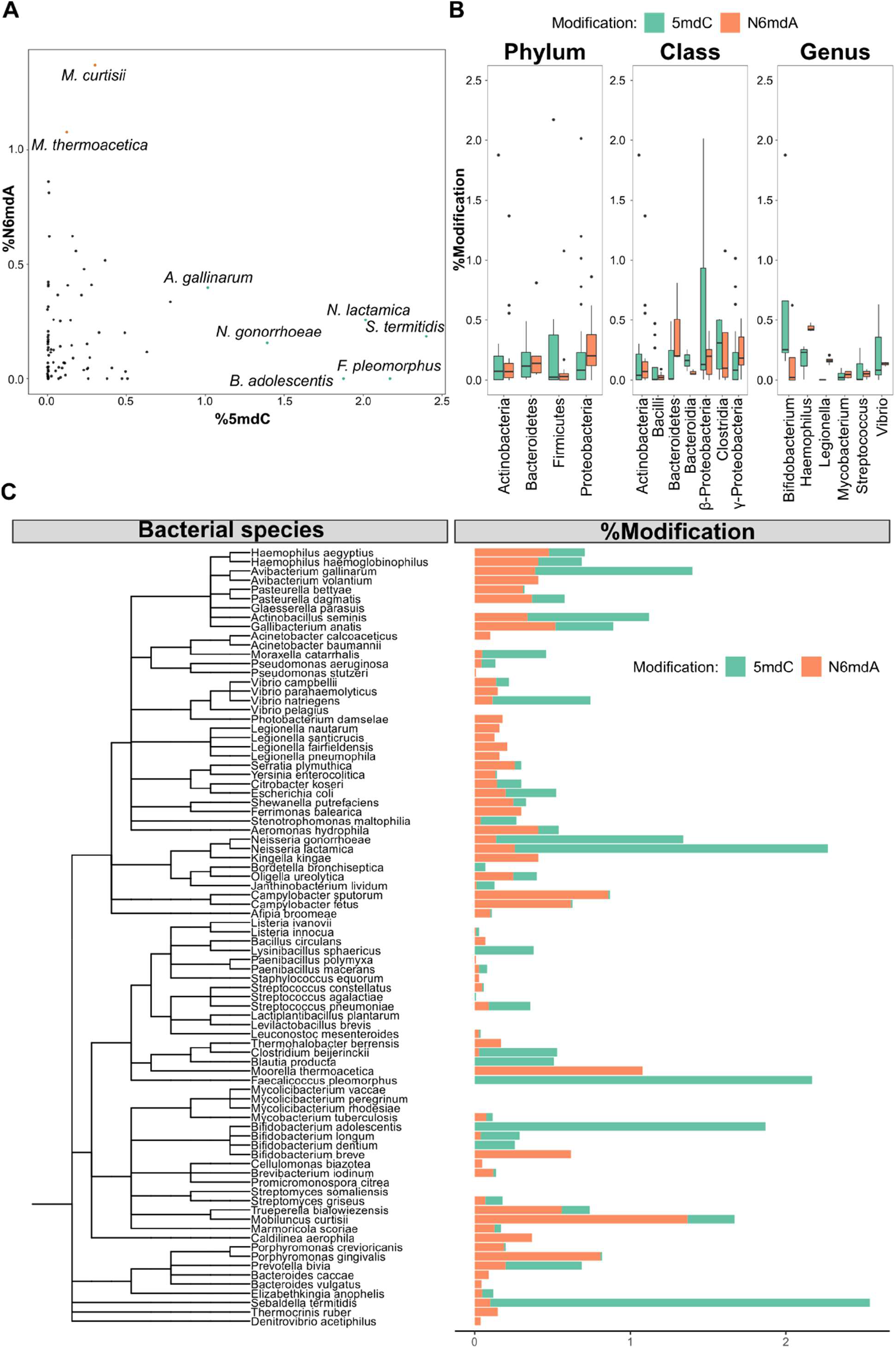
DNA modifications in bacteria. (A) Percentage of cytidine methylated against the percentage of adenine methylated in bacterial species. (B) Variation of % 5-methyl deoxycytidine and % N6-methyl deoxyadenosine among taxonomic divisions: phylum, class, and genus. (C) Distribution of 5mdC and N6mdA among 87 bacterial species measured against their phylogenetic diversity.

Finally, we also observed a third modification, 4mdC, to be frequent in prokaryotes. 4mdC occurred as an exclusive cytosine modification in *Legionella fairfieldensis, Bacteroides caccae, Acinetobacter baumannii, Listeria innocua, Bifidobacterium breve, Moorella thermoacetica, Thermocrinis ruber, Bacteroides vulgatus*, and *Caldilinea aerophila*. Moreover, it existed in tandem with 5mdC as a second modification in *Shewanella putrefaciens, Stenotrophomonas maltophilia, Bifidobacterium dentium, Mobiluncus curtisii*, and *Gallibacterium anatis* (not quantified).

Having analyzed 85 species, we were able to ask if bacterial species with a close evolutionary relationship or similar habitat or genome properties also have a more similar modification makeup. We did not detect any relationship between nature and level of modification and genome size or GC content (Fig. S5). Similarly, we detected no significant correlation between factors such as pathogenicity, temperature of growth, or tolerance to oxygen and the amount of modifications per unit genome size. We did however observe obvious patterns at the different taxonomic levels once we grouped the different bacterial strains according to phylum, class, and genus. Similarities are detected at the genus level (Fig. 4B–C). Members of the same genus often displayed similar values for a given modification. For example, species of the *Vibrio* genus presented similar quantities of N6mdA. At the class level we observed trends between the different classes and the amount of modification. α- and γ-Proteobacteria had the highest N6mdA content among different classes present while bacteroidetes presented with more cytidine methylation than adenosine methylation. At the phylum level the patterns were more prominent in Proteobacteria, containing more N6mdA than 5mdC, while a reverse trend of more 5mdC than N6mdA was observed for Bacteroidetes and Firmicutes. Combined, these results suggest that differences in the modifications do not reflect basic structural genome features such as size or GC content, but the observation that more-closely evolutionarily related species have higher similarities in DNA modification suggests that gene drift and gene function are key drivers in the evolution of DNA modifications.

## Acknowledgements

We thank Biological Research Facility at Francis Crick Institute for *Mus musculus, Danio rerio, Xenopus laevis*, samples, Bryony Lee (Turner lab, The Francis Crick Institute) for opossum samples, Annick Sawala (Gould Lab) for *Drosophila* samples, Cell Services (The Francis Crick Institute) for animal cell lines, National Yeast Collection for yeast samples, Felix Forest (Kew Gardens) and (Nell Jones) Chelsea Physic Garden for plant samples, Barbara Tautsher, Elisabeth Haring, Luise Kruckenhauser, for *Cepaea hortensis* and *Garra barremiae* samples (Natural History Museum of Vienna), Florian Winkler, Heinrich Aukenthaler, Erhard Seehauser, and Gottfried Hopfgartner (Forestry and Hunting Authorities South Tyrol, or Jagdrevier Mauls, Bolzano Province, Italy) for their support in obtaining tissue samples from alpine marmot in their wild habitats of Mauls and Gsies (Italy). This work was supported by the Francis Crick Institute which receives its core funding from Cancer Research UK (FC001134), the UK Medical Research Council (FC001134), and the Wellcome Trust (FC001134), and received specific support from the Wellcome Trust (200829/Z/16/Z, 101503/Z/13/Z) and the German Ministry of Education and Research (BMBF) as part of the National Research Node “Mass spectrometry in Systems Medicine (MSCoresys)”, under grant agreement 031L0220A.

