## SupplementaryInformation for "Occurrence and quantities of DNA modifications across the tree of life"

**Supplementary Information for**
**Occurrence and quantities of DNA modifications across the tree of life**

Sreejith Jayasree Varma[†1], Enrica Calvani[†2,3], Nana-Maria Grüning[1], Christoph Messner[2,3], Nicholas Grayson[5], Floriana Capuano[2], Michael Mülleder[4], Markus Ralser* [1,2,3]

[1] Department of Biochemistry, Charité - Universitätsmedizin Berlin, Germany.

[2] Department of Biochemistry and Cambridge Systems Biology Center, University of Cambridge, UK.

[3] The Molecular Biology of Metabolism Laboratory, The Francis Crick Institute, London, UK.

[4] Core Facility – High Throughput Mass Spectrometry, Charité - Universitätsmedizin Berlin, Germany.

[5] Wellcome Trust Sanger Institute, Wellcome Trust Genome Campus, Hinxton, United Kingdom.

[†]These authors contributed equally to the work
*To whom correspondence should be addressed. M. Ralser, Charité Universitätsmedizin Berlin.


Contents

1. Sample sources
2. Sample preparation
3. Data processing and analysis
4. Table of measured DNA modifications
5. Figures
6. References

1. **Sample sources**:
All the tissues, cell lines, yeast, and bacteria samples, unless differently stated, were extracted using Genomic Tip-20 kit (Qiagen) following the manufacturer's instructions.

Animals:

A. *Mus musculus*: DNA was extracted from spleen, brain, liver, heart, and lung of the following lines: FVB/NHanHsd, Balb/c, HsdOla:MF1, B6.SJL CD45.1/CD45.2, Balb/cAnN, 129S8, and F1(CBAxB6) of male individuals (8–11 weeks age).
The animals were provided by the Biological Research Facility at the Francis Crick Institute. Briefly, the animals were culled using rising concentrations of $CO_2$ over a cycle until it reached the 20%. The total duration of the procedure was 8 minutes. For death confirmation, before tissue dissection, the animals had their cervical vertebrae dislocated. After tissue dissection the samples were snap-frozen in liquid nitrogen and subsequently stored at –80 °C till further processing. All samples were obtained from the Biological Research Facility at the Francis Crick Institute.

A. *Monodelphis domestica*: DNA was extracted from the brain, liver, and kidney of 6 individuals kindly provided by the Turner lab at the Francis Crick Institute. The samples comprised of:
   - Three females aged 34 weeks and 6 days; 136 weeks; and 114 weeks and 3 days at the time of sacrifice.
   - Three male aged 68 weeks; 48 weeks and 4 days; and 46 weeks and 4 days at the time of sacrifice.

   Opossums were anesthetized by exposure to isoflurane gas until unresponsive to rocking of the cage and/or toe pinch. Subsequently, the animals were culled by cervical dislocation using a plastic scraper tool. Death was confirmed by the severing of the carotid artery.

B. *Danio rerio*: Seven 3rd-generation AB strain males (ZIRC) were sacrificed by an overdose of lidocaine (1g/L) buffered with sodium bicarbonate (2g/L) and subsequently frozen on dry ice. The samples were obtained by the Biological Research Facility at the Francis Crick Institute.

C. *Xenopus laevis*: Samples composed of tissues from the central nervous system (CNS), blood, heart, lung, liver, spleen, and kidney harvested from a male subject fertilized in March 2013. The samples were obtained by the Biological Research Facility at the Francis Crick Institute.

D. *Garra barreimiae*: DNA Samples were obtained from the Natural History Museum, Vienna, Austria.

E. *Marmota marmota*: Samples were harvested from four animals (two males, two females) used in a previous study (Gossmann et al. 2019). Each was obtained from three wild alpine marmot populations in the Central Alps near Mauls, Italy, at 2,367 m.a.s.l. at Mt Senges, 46° 52’ 40.55’’ N 11° 34’ 56.12’’ E, around St Martin, Gsies, Italy, at >2,000 m.a.s.l, 46° 49’ 44.2’’ N 12° 12’ 15.5’’ E, and in the nature reserve of La Grande Sassière (French Alps, 2,340 m.a.s.l., 45° 29’ N, 65° 90’ E).

F. *Cepaea hortensis*: Samples were obtained from the Natural History Museum, Vienna, Austria.

G. *Drosophila melanogaster*: Fly embryos were treated with 50% bleach to remove the microbiome around the cells and plated under following conditions:
One group of bleach-treated embryos were plated onto non-sterile food, while the other group was maintained microbiome free using sterile food. For comparison, non-bleached embryos were plated onto sterile and non-sterile food. In total there are therefore four treatment groups:

1) non-bleached embryos grown on non-sterile diet,
2) non-bleached embryos grown on sterile diet,
3) bleached embryos grown on non-sterile diet,
4) bleached embryos grown on sterile diet.

Upon hatching, groups of 15 male adult flies were collected and pooled to be considered as a biological replica and were processed using Tip-20.
(Annick Sawala, Gould Lab, the Francis Crick Institute)

Cell lines:
DNA was extracted from the following cell lines obtained from the STP Cell Services, the Francis Crick Institute: HeLa, Sf9, HCT116, MCF7, High 5 (2), 3t3 Balbc, CACO, HEK293, Hep G2, RAJI, HSB2, HBL-1, C13 (BHK-21), U937, Jurkat- E 6.1, THP-1, U20S, 293T. The number of cells processed was 5 x $10^6$ cells. The DNA was extracted using Genomic Tip-20 kit (Qiagen).

Bacteria:
The genomic DNA for species *Moorella thermoacetica, Halobacterium salinarum, Clostridium beijerinckii, Pseudomonas aeruginosa, Ferrimonas balearica, Vibrio parahaemolyticus, Yersinia enterocolitica sub palearctica, Yersinia enterocolitica sub enterocolitica, Denitrovibrio acetiphilus, Caldilinea aerophila, Marmoricola scoriae, Thermocrinis ruber, Streptococcus pneumoniae, Thermohalobacter berrensis, Promicromonospora citrea, Pseudomonas aeruginosa, Bacteroides vulgatus, Caldilinea aerophila,* and *Pseudomonas aeruginosa* was obtained from DSMZ as a solution in Tris/EDTA pH 8.0.

For the species *Avibacterium volantium, Serratia plymuthica, Sebaldella termitidis, Oligella ureolytica, Janthinobacterium lividum, Streptomyces griseus, Porphyromonas crevioricanis, Bordetella bronchiseptica, Acinetobacter calcoaceticus, Acinetobacter calcoaceticus, Porphyromonas gingivalis, Legionella fairfieldensis, Legionella santicrucis, Campylobacter sputorum biovar sputorum, Bacteroides caccae, Trueperella bialowiezensis, Neisseria gonorrhoeae, Afipia broomeae, Pasteurella bettyae, Escherichia coli, Janthinobacterium lividum, Acinetobacter baumannii, Brevibacterium iodinum, Citrobacter koseri, Shewanella putrefaciens, Listeria innocua, Legionella nautarum, Campylobacter fetus subsp. Fetus, Streptomyces somaliensis, Neisseria lactamica, Bacteroides vulgatus, Staphylococcus equorum, Bacillus circulans, Lactobacillus plantarum subsp. Plantarum, Escherichia coli, Streptococcus agalactiae, Legionella pneumophila subsp. Pascullei, Elizabethkingia anophelis, Cellulomonas biazotea, Pseudomonas stutzeri, Citrobacter koseri, Neisseria gonorrhoeae, Vibrio natriegens, Actinobacillus seminis, Lysinibacillus sphaericus, Mycobacterium rhodesiae, Stenotrophomonas maltophilia, Aeromonas hydrophila, Paenibacillus macerans, Paenibacillus polymyxa, Faecalicoccus pleomorphus, Streptococcus constellatus, Bifidobacterium breve, Bifidobacterium dentium, Mobiluncus curtisii, Haemophilus haemoglobinophilus, Haemophilus aegyptius, Haemophilus parasuis, Kingella kingae, Moraxella catarrhalis, Prevotella bivia, Vibrio pelagius, Avibacterium gallinarum, Vibrio campbellii, Vibrio natriegens, Neisseria gonorrhoeae, Pasteurella dagmatis, Mycobacterium peregrinum, Mycobacterium vaccae, Bifidobacterium longum subsp. Infantis, Blautia producta, Lactobacillus brevis, Listeria ivanovii subsp. Ivanovii, Gallibacterium anatis, Photobacterium damselae,* and *Bifidobacterium adolescentis* the genomic DNA was obtained from the Sanger Institute, Cambridge, UK. The DNA was extracted using MasterPure Complete DNA and RNA Purification Kit (Lucigen).

Yeast: Live cultures of *Wickerhamomyces anomalus, Zygosaccharomyces rouxii, Debaryomyces hansenii, Hanseniaspora osmophila, Rhodotorula mucilaginosa, Kluyveromyces marxianus, Lachancea thermotolerans,* and *Galactomyces candidus* were obtained from the National Collection of Yeast Cultures, Norwich, UK. The live cultures were grown on YPD agar overnight at room temperature, the colonies were then subcultured in YPD and incubated overnight at 25 °C. The cells were harvested at an $OD_{600}$ 1.0, corresponding roughly to $1.5 \times 10^9$ cells .

Plants:
Plant samples were obtained from the following sources:
*Helianthus annuus* as leaf tissue from Grüner Holländer Pflanzencenter Berlin;
*Capsicum annuum, Diplotaxis tenuifolia, Helianthus annuus* (seed), *Lepidium sativum* from seed from www.nebelung.de as seeds;
*Allium schoenoprasum* and *Salvia officinalis* from www.sperli.de as seeds;

*Spinacia oleracea, Cucurbita pepo, Phaseolus vulgaris* from www.Kiepenkerl.de as seeds;
*Artemisia dracunculus* from www.pflanzen-koelle.de as seeds;
*Andropogon gerardi* from www.plant-world-seeds.com as seeds;
*Raphanus sativus* from www.thompson-morgan.com as seeds;
*Cycas revoluta, Echinodorus grandiflorus* ssp *grandiflorus, Kleinia abyssinica* var *hildebrandtii, Sobralia macrantha* var *alba* were obtained from Kew Garden, London as leaf tissue.
*Ephedra viridis, Chamaerops humilis, Tetrastigma voinierianum, Nandina domestica, Viscum album, Magnolia grandiflora, Dracunculus canariensis* from Chelsea Physic Garden, London as leaf tissue.
Wild rocket from a local grocery store as leaf tissue.

For seed samples, the whole plant was harvested 7 days following germination at 25 °C on a water soaked tissue paper. The seeds were washed with 70% ethanol before preparing for germination. The obtained tissue was dried at 37 °C overnight, powdered using a mortar in liquid nitrogen and the DNA was extracted using a standard phenol-chloroform-isoamyl alcohol protocol (Allen et al. 2006).

2. **Sample preparation**

DNA extracts were treated with RNase A (VWR, Cat.No. E866-5ML) at 37 °C for 45 min and DNA purification was performed according to the manufacturer's instructions. Purified DNA was precipitated with isopropanol, washed with 70% ethanol, and resuspended in 10 mM Tris-HCl, pH 8.0. Quantification was done using a dsDNA BR Assay Kit (Qubit). The DNA sample was then digested into corresponding nucleosides using DNA Degradase Plus (Zymo Research, E2020). One µg of DNA was treated with 5 U of DNA Degradase at 37 °C for 2 h in a final volume of 25 µl and the reaction was inactivated, incubating the samples for 20 min at 70 °C as described by the manufacturer. Calibrations standards were prepared in 1:4:4:2:2:2:2:4:4:4:4:4:4:4:10 serial dilutions from a standard stock that was prepared according to the following:

**Table S1:** Concentrations of pure nucleoside standards and their sources

| **Molecule: Vendor/Code** | **Pure Stock Concentration µM** | **Pool concentration µM** |
|---|---|---|
| 2dC: Sigma/D3897-100MG | 5000 | 100 |
| 5hmdC: Berry and Associates/PY7588 | 0.5 | 0.04 |
| 5mdC: Santa Cruz/ sc-278256 | 100 | 0.02 |
| cadC: Berry and Associates/PY7593 | 0.5 | 0.02 |

| | | |
|---|---|---|
| dA: Sigma/D7400-250MG | 5000 | 100 |
| dG: Sigma/854999 | 5000 | 100 |
| fdC: Berry and Associates/PY 7589 | 0.5 | 0.02 |
| N6mdA: Alfa Aesar/ J64961 | 0.5 | 0.02 |
| T: Sigma/89270-1G | 5000 | 100 |

The samples were diluted 1:1 with MeOH 10% (v/v) containing 0.2% formic acid, and 10 µl corresponding to 200 ng of gDNA were injected onto a reverse phase chromatography Acquity UPLC HSS T3 column, 100 Å, 1.8 µm, 2.1 mm x150 mm (Waters), column temperature 25 °C and flow rate of 0.2 mL/min. Mobile phase A: 0.1% formic acid + 10 mM ammonium formate in water, mobile phase B: 0.1% formic acid + 10 mM ammonium formate in methanol. Gradient for elution was started from 5% mobile phase B to 35% B over 11.5 minutes followed by sharply increasing to 80% over the next 1.5 minutes. The gradient was held at 80% B for 2 minutes, lowered to the starting gradient over 1 minute and equilibrated for 6.5 minutes. Total length was 22.5 minutes.
The eluent was directed to an electrospray ion source connected to a triple quadrupole mass spectrometer (Agilent 6460 QQQ) equipped with an Agilent Jet stream source, operating in positive mode. The ESI source settings were: gas temperature: 300 °C; gas flow: 6.4 L/min; nebulizer: 50 psi; sheath gas heater: 350 °C; sheath gas flow: 7 L/min; capillary: 2000 V. The following transitions were monitored for MRM experiments:

**Table S2:** Retention times and transitions for nucleosides analyzed

| **Molecule** | **Precursor ion** | **Qualifier Product ion** | **Quantifier Product ion** | **Retention time (min)** | **Mode** |
|---|---|---|---|---|---|
| 2dC | 228.1 | 95 | 112.0 | 4.362 | Positive |
| 5hmdC | 258.0 | 141.9 | 81.1 | 4.342 | Positive |
| 5mdC | 242.0 | 108.6 | 126.0 | 5.655 | Positive |
| cadC | 272.0 | 137.9 | 155.9 | 5.193 | Positive |
| dA | 252.1 | - | 136.0 | 8.128 | Positive |
| dG | 268.1 | - | 151.9 | 7.546 | Positive |
| fdC | 256.0 | 97 | 139.9 | 7.868 | Positive |
| N6mdA | 266.3 | 117 | 150.0 | 11.391 | Positive |

| | | | | | |
|---|---|---|---|---|---|
| T | 243.1 | 54.1 | 126.9 | 8.349 | Positive |

For Neutral loss measurements, the samples were injected as per the same LC parameters used for the MRM experiment while the mass spectrometer was set to a scan type of Neutral loss (M = 116 Da) while scanning the quadrupoles from 230 to 250 Da. The scan time was 1000 with step size of 0.05 amu and the values for Fragmentor, collision energy and cell accelerator voltage were 73, 8 and 5 respectively.

3. **Data processing and analysis**

Peak areas were extracted and integrated using MassHunter for QQQ to obtain the concentrations after applying the necessary limits of quantification. Subsequent processing for batch-to-batch variation and technical outlier removal were carried out using R or Python. A single reference mouse DNA sample was included in every measured batch to monitor batch-to-batch variation. Median-value-based normalization of the reference mouse samples was used to obtain the correction factor with which the corresponding batch was corrected. The results are depicted as percentage modification with respect to dG (for 5mdC and 5hmdC) and T (for N6mdA). The phylogenetic clustering was carried out using a newick file generated using NCBI Taxonomy (PhyloT) and the ggtree package (Yu 2020). Features of bacteria were retrieved from the bacterial metadatabase BacDive (http://bacdive.dsmz.de, accessed 14 April, 2020).(Reimer et al. 2019)

4. **Table of measured DNA modifications**

The values are presented as % modification: %5mdC as (5mdC/dG)*100, %5hmdC as (5hmdC/dG)*100 and %N6mdA as (N6mdA/T)*100.

**Table S3:** Percentages of 5mdC (wrt dG), 5hmdC (wrt dG), and N6mdA (wrt T) of different samples measured*

| | **Species** | **Details** | **5mdC** | **5hmdC** | **N6mdA** | **Kingdom** | **Phylum** |
|---|---|---|---|---|---|---|---|
| 1 | *Acinetobacter baumannii* | | 0.33 | 0.00 | 0.02 | Monera | Proteobacteria |
| 2 | *Acinetobacter calcoaceticus* | | 0.01 | 0.00 | 0.37 | Monera | Proteobacteria |
| 3 | *Acinetobacter calcoaceticus* | | 0.01 | 0.00 | 0.11 | Monera | Proteobacteria |
| 4 | *Actinobacillus seminis* | | 1.88 | 0.00 | 0.81 | Monera | Proteobacteria |

| | | | | | | | |
|---|---|---|---|---|---|---|---|
| 5 | *Aeromonas hydrophila* | | 0.63 | 0.00 | 2.01 | Monera | Proteobacteria |
| 6 | *Afipia broomeae* | | 0.05 | 0.00 | 0.67 | Monera | Proteobacteria |
| 7 | *Allium cepa* | whole seedling | 37.12 | 0.09 | 0.01 | Plantae | Tracheophyta |
| 8 | *Allium schoenoprasum* | whole seedling | 25.67 | 0.00 | 0.04 | Plantae | Tracheophyta |
| 9 | *Andropogon gerardi* | whole seedling | 37.51 | 0.00 | 0.00 | Plantae | Tracheophyta |
| 10 | *Artemisia dracunculus* | whole seedling | 23.22 | 0.00 | 0.00 | Plantae | Tracheophyta |
| 11 | *Avibacterium gallinarum* | | 2.38 | 0.00 | 0.93 | Monera | Proteobacteria |
| 12 | *Avibacterium volantium* | | 0.01 | 0.00 | 0.90 | Monera | Proteobacteria |
| 13 | *Bacillus circulans* | | 0.01 | 0.00 | 0.35 | Monera | Firmicutes |
| 14 | *Bacteroides caccae* | | 0.88 | 0.00 | 0.48 | Monera | Bacteriodetes |
| 15 | *Bacteroides vulgatus* | DSM 1447 | 0.03 | 0.00 | 0.15 | Monera | Bacteriodetes |
| 16 | *Bifidobacterium adolescentis* | | 4.13 | 0.00 | 0.00 | Monera | Actinobacteria |
| 17 | *Bifidobacterium breve* | | 0.37 | 0.00 | 1.45 | Monera | Actinobacteria |
| 18 | *Bifidobacterium dentium* | | 0.68 | 0.00 | 0.00 | Monera | Actinobacteria |
| 19 | *Bifidobacterium longum* | subsp. *infantis* | 0.60 | 0.00 | 0.10 | Monera | Actinobacteria |
| 20 | *Blautia producta* | | 3.09 | 0.00 | 0.00 | Monera | Firmicutes |
| 21 | *Bordetella bronchiseptica* | | 0.38 | 0.00 | 0.00 | Monera | Proteobacteria |
| 22 | *Brevibacterium iodinum* | | 0.07 | 0.00 | 0.41 | Monera | Actinobacteria |

| | | | | | | | |
|---|---|---|---|---|---|---|---|
| 23 | *Caldilinea aerophila* | DSM 14535 | 0.34 | 0.00 | 1.88 | Monera | Chloroflexi |
| 24 | *Campylobacter fetus* | subsp. *fetus* | 0.02 | 0.00 | 1.13 | Monera | Proteobacteria |
| 25 | *Campylobacter sputorum* | biovar *sputorum* | 0.01 | 0.00 | 1.51 | Monera | Proteobacteria |
| 26 | *Candida tropicalis* | | 0.02 | 0.00 | 0.00 | Fungi | Ascomycota |
| 27 | *Capsicum annuum* | whole seedling | 29.57 | 0.00 | 0.02 | Plantae | Tracheophyta |
| 28 | *Cellulomonas biazotea* | | 0.00 | 0.00 | 0.22 | Monera | Actinobacteria |
| 29 | *Cepaea hortensis* | Gland | 0.05 | 0.00 | 0.00 | Animalia | Mollusca |
| 30 | *Cepaea hortensis* | Muscle | 0.84 | 0.32 | 0.01 | Animalia | Mollusca |
| 31 | *Citrobacter koseri* | | 1.47 | 0.00 | 1.34 | Monera | Proteobacteria |
| 32 | *Citrobacter koseri* | | 0.01 | 0.00 | 0.06 | Monera | Proteobacteria |
| 33 | *Clostridium beijerinckii* | DSM 791 | 2.97 | 0.00 | 0.18 | Monera | Firmicutes |
| 34 | *Cucurbita maxima* | whole seedling | 11.32 | 0.00 | 0.03 | Plantae | Tracheophyta |
| 35 | *Cucurbita pepo* | whole seedling | 16.61 | 0.00 | 0.01 | Plantae | Tracheophyta |
| 36 | *Cupressus torulosa* | whole seedling | 22.22 | 0.00 | 0.01 | Plantae | Tracheophyta |
| 37 | *Danio rerio* | whole body | 10.57 | 0.25 | 0.00 | Animalia | Chordata |
| 38 | *Debaryomyces hansenii* | | 0.02 | 0.00 | 0.00 | Fungi | Ascomycota |
| 39 | *Denitrovibrio acetiphilus* | DSM 12809 | 0.01 | 0.00 | 0.13 | Monera | Deferribacteres |
| 40 | *Diplotaxis tenuifolia* | whole seedling | 21.26 | 0.00 | 0.05 | Plantae | Tracheophyta |

| | | | | | | | |
|---|---|---|---|---|---|---|---|
| 41 | *Drosophila melanogaster* | Male/Bleached Nonsterile | 0.00 | 0.00 | 0.02 | Animalia | Arthropoda |
| 42 | *Drosophila melanogaster* | Male/Bleached Sterile | 0.00 | 0.00 | 0.03 | Animalia | Arthropoda |
| 43 | *Drosophila melanogaster* | Male/Unbleached Nonsterile | 0.00 | 0.09 | 0.06 | Animalia | Arthropoda |
| 44 | *Drosophila melanogaster* | Male/Unbleached Sterile | 0.00 | 0.00 | 0.04 | Animalia | Arthropoda |
| 45 | *Elizabethkingia anophelis* | | 0.29 | 0.00 | 0.20 | Monera | Bacteriodetes |
| 46 | *Escherichia coli* | | 1.70 | 0.00 | 1.00 | Monera | Proteobacteria |
| 47 | *Escherichia coli* | | 1.65 | 0.00 | 1.07 | Monera | Proteobacteria |
| 48 | *Faecalicoccus pleomorphus* | | 4.46 | 0.00 | 0.00 | Monera | Firmicutes |
| 49 | *Ferrimonas balearica* | DSM 9799 | 0.00 | 0.00 | 1.29 | Monera | Proteobacteria |
| 50 | *Gallibacterium anatis* | | 0.91 | 0.00 | 1.29 | Monera | Proteobacteria |
| 51 | *Garra barreimiae* | Surface muscle | 5.76 | 1.09 | 0.00 | Animalia | Chordata |
| 52 | *Garra barreimiae* | Surface CNS | 6.44 | 8.83 | 0.00 | Animalia | Chordata |
| 53 | *Garra barreimiae* | Cave CNS | 6.01 | 8.29 | 0.00 | Animalia | Chordata |
| 54 | *Garra barreimiae* | Cave muscle | 5.81 | 1.00 | 0.00 | Animalia | Chordata |
| 55 | *Geotrichum candidum* | | 0.27 | 0.00 | 0.00 | Fungi | Ascomycota |
| 56 | *Haemophilus aegyptius* | | 0.45 | 0.00 | 0.93 | Monera | Proteobacteria |
| 57 | *Haemophilus haemoglobinophilus* | | 0.67 | 0.00 | 0.98 | Monera | Proteobacteria |

| | | | | | | | |
|---|---|---|---|---|---|---|---|
| 58 | *Haemophilus parasuis* | | 0.01 | 0.00 | 0.95 | Monera | Proteobacteria |
| 59 | *Halobacterium salinarum* | DSM 670 | 0.03 | 0.01 | 0.10 | Monera | Euryarchaeota |
| 60 | *Hanseniaspora osmophila* | | 0.04 | 0.00 | 0.00 | Fungi | Ascomycota |
| 61 | *Helianthus annuus* | Leaf | 35.22 | 0.00 | 0.00 | Plantae | Tracheophyta |
| 62 | *Homo sapiens* | Leukemic monocyte cancer cells | 3.08 | 0.00 | 0.00 | Animalia | Chordata |
| 63 | *Homo sapiens* | Burkitt's lymphoma cancer cells | 3.14 | 0.00 | 0.00 | Animalia | Chordata |
| 64 | *Homo sapiens* | Cervical cancer cells | 1.14 | 0.00 | 0.00 | Animalia | Chordata |
| 65 | *Homo sapiens* | Colon adenocarcinoma cancer cells | 1.56 | 0.00 | 0.00 | Animalia | Chordata |
| 66 | *Homo sapiens* | Colon cancer cells | 1.51 | 0.00 | 0.00 | Animalia | Chordata |
| 67 | *Homo sapiens* | Embryo kidney cells | 1.45 | 0.00 | 0.00 | Animalia | Chordata |
| 68 | *Homo sapiens* | Hepatocyte carcinoma cancer cells | 1.49 | 0.00 | 0.00 | Animalia | Chordata |
| 69 | *Homo sapiens* | Lymphoblast | 5.38 | 0.00 | 0.01 | Animalia | Chordata |
| 70 | *Homo sapiens* | Osteosarcoma cancer cells | 2.43 | 0.00 | 0.00 | Animalia | Chordata |
| 71 | *Homo sapiens* | T cells | 4.92 | 0.00 | 0.00 | Animalia | Chordata |
| 72 | *Homo sapiens* | T cell leukemia cancer cells | 2.95 | 0.00 | 0.00 | Animalia | Chordata |
| 73 | *Homo sapiens* | Breast cancer cells | 1.36 | 0.01 | 0.00 | Animalia | Chordata |

| | | | | | | | |
|---|---|---|---|---|---|---|---|
| 74 | *Homo sapiens* | B cell lymphoma cancer | 4.66 | 0.00 | 0.01 | Animalia | Chordata |
| 75 | *Janthinobacterium lividum* | | 0.01 | 0.00 | 0.11 | Monera | Proteobacteria |
| 76 | *Janthinobacterium lividum* | | 1.46 | 0.00 | 0.08 | Monera | Proteobacteria |
| 77 | *Kingella kingae* | | 0.01 | 0.00 | 0.82 | Monera | Proteobacteria |
| 78 | *Kluyveromyces marxianus* | | 0.02 | 0.00 | 0.00 | Fungi | Ascomycota |
| 79 | *Lachancea thermotolerans* | | 0.02 | 0.00 | 0.00 | Fungi | Ascomycota |
| 80 | *Lactobacillus brevis* | | 1.21 | 0.00 | 0.00 | Monera | Firmicutes |
| 81 | *Lactobacillus plantarum* | subsp. *plantarum* | 0.02 | 0.00 | 0.00 | Monera | Firmicutes |
| 82 | *Laurus nobilis* | Leaf | 25.74 | 0.47 | 0.00 | Plantae | Tracheophyta |
| 83 | *Legionella fairfieldensis* | | 0.01 | 0.00 | 0.55 | Monera | Proteobacteria |
| 84 | *Legionella nautarum* | | 0.01 | 0.00 | 0.59 | Monera | Proteobacteria |
| 85 | *Legionella pneumophila* | subsp. pascullei | 0.01 | 0.00 | 0.55 | Monera | Proteobacteria |
| 86 | *Legionella santicrucis* | | 0.01 | 0.00 | 0.61 | Monera | Proteobacteria |
| 87 | *Lepidium sativum* | whole seedling | 10.53 | 1.67 | 0.00 | Plantae | Tracheophyta |
| 88 | *Leuconostoc mesenteroides* | | 0.01 | 0.00 | 0.06 | Monera | Firmicutes |
| 89 | *Listeria innocua* | | 0.07 | 0.00 | 0.03 | Monera | Firmicutes |
| 90 | *Listeria ivanovii* | subsp. *ivanovii* | 0.01 | 0.00 | 0.00 | Monera | Firmicutes |

| | | | | | | | |
|---|---|---|---|---|---|---|---|
| 91 | *Lysinibacillus sphaericus* | | 1.75 | 0.00 | 0.00 | Monera | Firmicutes |
| 92 | *Marmoricola scoriae* | DSM 22127 | 0.16 | 0.00 | 0.51 | Monera | Actinobacteria |
| 93 | *Marmota marmota* | Kidney | 5.08 | 0.32 | 0.00 | Animalia | Chordata |
| 94 | *Marmota marmota* | Liver | 2.86 | 0.38 | 0.00 | Animalia | Chordata |
| 95 | *Marmota marmota* | Spleen | 4.89 | 0.08 | 0.00 | Animalia | Chordata |
| 96 | *Mesocricetus auratus* | not available | 1.44 | 0.00 | 0.00 | Animalia | Chordata |
| 97 | *Mimosa pudica* | Leaf | 10.15 | 0.00 | 0.00 | Plantae | Tracheophyta |
| 98 | *Mobiluncus curtisii* | | 0.65 | 0.00 | 2.95 | Monera | Actinobacteria |
| 99 | *Monodelphis domestica* | Female liver | 1.81 | 1.36 | 0.00 | Animalia | Chordata |
| 100 | *Monodelphis domestica* | Male CNS | 2.24 | 0.42 | 0.00 | Animalia | Chordata |
| 101 | *Monodelphis domestica* | Male kidney | 2.00 | 0.41 | 0.00 | Animalia | Chordata |
| 102 | *Monodelphis domestica* | Male liver | 1.87 | 1.52 | 0.00 | Animalia | Chordata |
| 103 | *Monodelphis domestica* | Female kidney | 1.84 | 0.79 | 0.00 | Animalia | Chordata |
| 104 | *Monodelphis domestica* | Female CNS | 2.11 | 1.05 | 0.00 | Animalia | Chordata |
| 105 | *Moorella thermoacetica* | DSM 521 | 0.32 | 0.00 | 2.83 | Monera | Firmicutes |
| 106 | *Moraxella catarrhalis* | | 0.78 | 0.01 | 0.09 | Monera | Proteobacteria |
| 107 | *Mus musculus* | FVB/N lung | 4.73 | 0.48 | 0.00 | Animalia | Chordata |
| 108 | *Mus musculus* | FVB/N liver | 4.88 | 0.55 | 0.00 | Animalia | Chordata |

| | | | | | | | |
|---|---|---|---|---|---|---|---|
| 109 | *Mus musculus* | Hsd/Ola/MF1 lung | 4.68 | 0.57 | 0.00 | Animalia | Chordata |
| 110 | *Mus musculus* | Hsd/Ola/MF1 brain | 5.40 | 3.36 | 0.00 | Animalia | Chordata |
| 111 | *Mus musculus* | Hsd/Ola/MF1 heart | 4.32 | 0.59 | 0.00 | Animalia | Chordata |
| 112 | *Mus musculus* | Hsd/Ola/MF1 liver | 4.79 | 0.55 | 0.00 | Animalia | Chordata |
| 113 | *Mus musculus* | FVB/N heart | 3.85 | 0.56 | 0.00 | Animalia | Chordata |
| 114 | *Mus musculus* | Hsd/Ola/MF1 spleen | 5.06 | 0.04 | 0.00 | Animalia | Chordata |
| 115 | *Mus musculus* | FVB/N spleen | 4.85 | 0.06 | 0.00 | Animalia | Chordata |
| 116 | *Mus musculus* | FVB/N brain | 5.33 | 3.23 | 0.00 | Animalia | Chordata |
| 117 | *Mus musculus* | BALB/cAnN lung | 4.49 | 0.49 | 0.00 | Animalia | Chordata |
| 118 | *Mus musculus* | F1/CBAxB6 lung | 4.80 | 0.42 | 0.00 | Animalia | Chordata |
| 119 | *Mus musculus* | BALB/cAnN liver | 5.03 | 0.43 | 0.00 | Animalia | Chordata |
| 120 | *Mus musculus* | BALB/cAnN heart | 3.98 | 1.61 | 0.00 | Animalia | Chordata |
| 121 | *Mus musculus* | BALB/cAnN brain | 5.55 | 3.45 | 0.00 | Animalia | Chordata |
| 122 | *Mus musculus* | BALB/c brain | 5.42 | 3.55 | 0.00 | Animalia | Chordata |
| 123 | *Mus musculus* | B6SJL/CD451/CD452 spleen | 4.76 | 0.04 | 0.00 | Animalia | Chordata |
| 124 | *Mus musculus* | B6SJL/CD451/CD452 lung | 4.47 | 0.39 | 0.00 | Animalia | Chordata |
| 125 | *Mus musculus* | B6SJL/CD451/CD452 liver | 4.80 | 0.43 | 0.00 | Animalia | Chordata |

| | | | | | | | |
|---|---|---|---|---|---|---|---|
| 126 | *Mus musculus* | B6SJL/CD451/CD452 heart | 4.58 | 0.58 | 0.00 | Animalia | Chordata |
| 127 | *Mus musculus* | B6SJL/CD451/CD452 brain | 5.30 | 3.19 | 0.00 | Animalia | Chordata |
| 128 | *Mus musculus* | BALB/c embryonic cells | 4.62 | 0.00 | 0.02 | Animalia | Chordata |
| 129 | *Mus musculus* | 129S8 spleen | 4.94 | 0.08 | 0.00 | Animalia | Chordata |
| 130 | *Mus musculus* | 129S8 lung | 4.89 | 0.55 | 0.00 | Animalia | Chordata |
| 131 | *Mus musculus* | 129S8 liver | 4.08 | 0.43 | 0.00 | Animalia | Chordata |
| 132 | *Mus musculus* | 129S8 heart | 4.54 | 0.49 | 0.00 | Animalia | Chordata |
| 133 | *Mus musculus* | 129S8 brain | 5.04 | 2.85 | 0.00 | Animalia | Chordata |
| 134 | *Mus musculus* | BALB/c heart | 4.90 | 0.48 | 0.00 | Animalia | Chordata |
| 135 | *Mus musculus* | BALB/c liver | 4.92 | 0.55 | 0.00 | Animalia | Chordata |
| 136 | *Mus musculus* | BALB/cAnN spleen | 4.69 | 0.06 | 0.00 | Animalia | Chordata |
| 137 | *Mus musculus* | BALB/c spleen | 4.82 | 0.07 | 0.00 | Animalia | Chordata |
| 138 | *Mus musculus* | F1/CBAxB6 brain | 5.40 | 3.43 | 0.00 | Animalia | Chordata |
| 139 | *Mus musculus* | F1/CBAxB6 heart | 5.79 | 0.88 | 0.00 | Animalia | Chordata |
| 140 | *Mus musculus* | F1/CBAxB6 liver | 4.98 | 0.38 | 0.00 | Animalia | Chordata |
| 141 | *Mus musculus* | F1/CBAxB6 spleen | 4.96 | 0.06 | 0.00 | Animalia | Chordata |
| 142 | *Mus musculus* | BALB/c lung | 4.64 | 0.50 | 0.00 | Animalia | Chordata |

| | | | | | | | |
|---|---|---|---|---|---|---|---|
| 143 | *Mycobacterium peregrinum* | | 0.70 | 0.00 | 0.47 | Monera | Actinobacteria |
| 144 | *Mycobacterium rhodesiae* | | 0.01 | 0.00 | 0.09 | Monera | Actinobacteria |
| 145 | *Mycobacterium tuberculosis* | | 0.06 | 0.00 | 0.13 | Monera | Actinobacteria |
| 146 | *Mycobacterium tuberculosis* | | 0.24 | 0.00 | 0.48 | Monera | Actinobacteria |
| 147 | *Mycobacterium tuberculosis* | | 0.17 | 0.00 | 0.16 | Monera | Actinobacteria |
| 148 | *Mycobacterium tuberculosis* | | 0.18 | 0.00 | 0.47 | Monera | Actinobacteria |
| 149 | *Mycobacterium vaccae* | | 0.00 | 0.00 | 0.13 | Monera | Actinobacteria |
| 150 | *Neisseria gonorrhoeae* | | 3.26 | 0.00 | 0.30 | Monera | Proteobacteria |
| 151 | *Neisseria gonorrhoeae* | | 2.41 | 0.00 | 0.19 | Monera | Proteobacteria |
| 152 | *Neisseria gonorrhoeae* | | 2.59 | 0.00 | 0.15 | Monera | Proteobacteria |
| 153 | *Neisseria gonorrhoeae* | | 2.33 | 0.00 | 0.36 | Monera | Proteobacteria |
| 154 | *Neisseria gonorrhoeae* | | 2.51 | 0.00 | 0.21 | Monera | Proteobacteria |
| 155 | *Neisseria gonorrhoeae* | | 2.45 | 0.00 | 0.59 | Monera | Proteobacteria |
| 156 | *Neisseria gonorrhoeae* | | 5.68 | 0.00 | 0.63 | Monera | Proteobacteria |
| 157 | *Neisseria gonorrhoeae* | | 3.09 | 0.00 | 0.14 | Monera | Proteobacteria |
| 158 | *Neisseria gonorrhoeae* | | 2.57 | 0.00 | 0.44 | Monera | Proteobacteria |
| 159 | *Neisseria lactamica* | | 4.40 | 0.00 | 0.56 | Monera | Proteobacteria |

| | | | | | | | |
|---|---|---|---|---|---|---|---|
| 160 | *Oligella ureolytica* | | 0.39 | 0.00 | 0.68 | Monera | Proteobacteria |
| 161 | *Paenibacillus macerans* | | 0.39 | 0.00 | 0.24 | Monera | Firmicutes |
| 162 | *Paenibacillus polymyxa* | | 0.01 | 0.00 | 0.05 | Monera | Firmicutes |
| 163 | *Pasteurella bettyae* | | 0.01 | 0.00 | 0.72 | Monera | Proteobacteria |
| 164 | *Pasteurella dagmatis* | | 0.49 | 0.00 | 0.83 | Monera | Proteobacteria |
| 165 | *Phaseolus vulgaris* | Leaf | 17.76 | 0.00 | 0.00 | Plantae | Tracheophyta |
| 166 | *Phaseolus vulgaris* | Root | 13.94 | 0.00 | 0.10 | Plantae | Tracheophyta |
| 167 | *Phaseolus vulgaris* | Stem | 16.69 | 0.00 | 0.01 | Plantae | Tracheophyta |
| 168 | *Phaseolus vulgaris* | empty cotyledon | 16.62 | 0.00 | 0.00 | Plantae | Tracheophyta |
| 169 | *Phaseolus vulgaris* | whole seedling | 18.78 | 0.00 | 0.01 | Plantae | Tracheophyta |
| 170 | *Photobacterium damselae* | | 0.01 | 0.00 | 0.81 | Monera | Actinobacteria |
| 171 | *Plasmodium falciparum* | | 0.11 | 0.00 | 0.00 | Protozoa | Apicomplexa |
| 172 | *Porphyromonas crevioricanis* | | 0.01 | 0.00 | 0.39 | Monera | Bacteriodetes |
| 173 | *Porphyromonas gingivalis* | | 0.02 | 0.00 | 1.82 | Monera | Bacteriodetes |
| 174 | *Prevotella bivia* | | 1.22 | 0.00 | 0.50 | Monera | Bacteriodetes |
| 175 | *Promicromonospora citrea* | DSM 46089 | 0.01 | 0.00 | 0.68 | Monera | Actinobacteria |
| 176 | *Pseudomonas aeruginosa* | DSM 50071 | 1.14 | 0.00 | 0.06 | Monera | Proteobacteria |

| | | | | | | | |
|---|---|---|---|---|---|---|---|
| 177 | *Pseudomonas aeruginosa* | DSM 1117 | 0.09 | 0.00 | 0.56 | Monera | Proteobacteria |
| 178 | *Pseudomonas stutzeri* | | 0.01 | 0.00 | 0.04 | Monera | Proteobacteria |
| 179 | *Raphanus sativus* | whole seedling | 3.43 | 0.14 | 0.01 | Plantae | Tracheophyta |
| 180 | *Salvia officinalis* | whole seedling | 13.17 | 0.00 | 0.00 | Plantae | Tracheophyta |
| 181 | *Sebaldella termitidis* | | 0.13 | 0.00 | 0.01 | Monera | Fusobacteria |
| 182 | *Serratia plymuthica* | | 0.21 | 0.00 | 1.40 | Monera | Proteobacteria |
| 183 | *Shewanella putrefaciens* | | 0.38 | 0.00 | 1.17 | Monera | Proteobacteria |
| 184 | *Sorghum bicolor* | whole seedling | 31.53 | 0.00 | 0.03 | Plantae | Tracheophyta |
| 185 | *Spinacia oleracea* | whole seedling | 26.51 | 0.00 | 0.04 | Plantae | Tracheophyta |
| 186 | *Spodoptera frugiperda* | Sf21 insect cells | 0.12 | 0.00 | 0.00 | Animalia | Arthropoda |
| 187 | *Staphylococcus equorum* | | 0.01 | 0.00 | 0.09 | Monera | Firmicutes |
| 188 | *Stenotrophomonas maltophilia* | | 1.05 | 0.00 | 0.17 | Monera | Proteobacteria |
| 189 | *Streptococcus agalactiae* | | 0.01 | 0.00 | 0.01 | Monera | Firmicutes |
| 190 | *Streptococcus constellatus* | | 0.01 | 0.00 | 0.10 | Monera | Firmicutes |
| 191 | *Streptococcus pneumoniae* | DSM 24048 | 0.56 | 0.00 | 0.19 | Monera | Firmicutes |
| 192 | *Streptomyces griseus* | | 0.79 | 0.00 | 0.49 | Monera | Actinobacteria |
| 193 | *Streptomyces somaliensis* | | 0.02 | 0.00 | 0.00 | Monera | Actinobacteria |

| | | | | | | | |
|---|---|---|---|---|---|---|---|
| 194 | *Sulfolobus acidocaldarius* | DSM 639 | 0.32 | 0.00 | 0.00 | Monera | Crenarchaeota |
| 195 | *Thermocrinis ruber* | DSM 23557 | 0.38 | 0.00 | 0.23 | Monera | Aquificae |
| 196 | *Thermohalobacter berrensis* | DSM 26700 | 0.01 | 0.00 | 0.47 | Monera | Firmicutes |
| 197 | *Trichoplusia ni* | Ovarian cells | 0.21 | 0.00 | 0.00 | Animalia | Arthropoda |
| 198 | *Trueperella bialowiezensis* | | 0.37 | 0.00 | 1.14 | Monera | Actinobacteria |
| 199 | *Valerianella locusta* | whole seedling | 3.32 | 0.00 | 0.01 | Plantae | Tracheophyta |
| 200 | *Vibrio campbellii* | | 0.45 | 0.00 | 0.78 | Monera | Proteobacteria |
| 201 | *Vibrio natriegens* | | 6.51 | 0.00 | 0.44 | Monera | Proteobacteria |
| 202 | *Vibrio natriegens* | | 0.00 | 0.00 | 0.76 | Monera | Proteobacteria |
| 203 | *Vibrio parahaemolyticus* | DSM 10027 | 0.01 | 0.00 | 0.77 | Monera | Proteobacteria |
| 204 | *Vibrio pelagius* | | 0.01 | 0.01 | 0.89 | Monera | Proteobacteria |
| 205 | *Xenopus laevis* | CNS | 7.33 | 2.00 | 0.00 | Animalia | Chordata |
| 206 | *Xenopus laevis* | Heart | 9.60 | 0.02 | 0.00 | Animalia | Chordata |
| 207 | *Xenopus laevis* | Kidney | 9.53 | 0.25 | 0.00 | Animalia | Chordata |
| 208 | *Xenopus laevis* | Liver | 10.25 | 0.04 | 0.00 | Animalia | Chordata |
| 209 | *Xenopus laevis* | Lung | 10.00 | 0.10 | 0.00 | Animalia | Chordata |
| 210 | *Xenopus laevis* | Spleen | 9.76 | 0.05 | 0.00 | Animalia | Chordata |

| | | | | | | | |
|---|---|---|---|---|---|---|---|
| 211 | *Yersinia enterocolitica* | sub enterocolitica-DSM 9499 | 0.10 | 0.00 | 0.58 | Monera | Proteobacteria |
| 212 | *Yersinia enterocolitica* | sub palearctica-DSM 11502 | 0.02 | 0.00 | 0.62 | Monera | Proteobacteria |

* Percentages are <u>NOT</u> normalized to genome size.

### 5. Figures

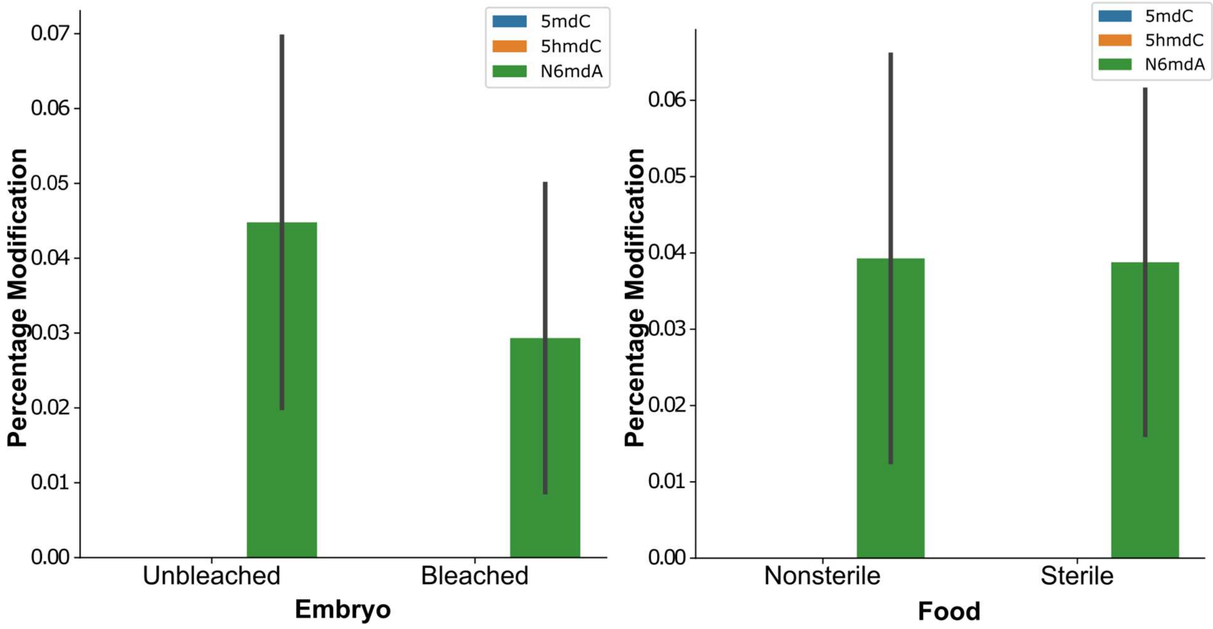


**Figure S1:** The variation of the three modifications 5mdC, 5hmdC, and N6mdA in *Drosophila melanogaster* in (A) bleached (microbiota removed via bleaching) vs. unbleached (intact microbiota) flies and in (B) bleached flies fed with a sterilized vs. non-sterilized diet.

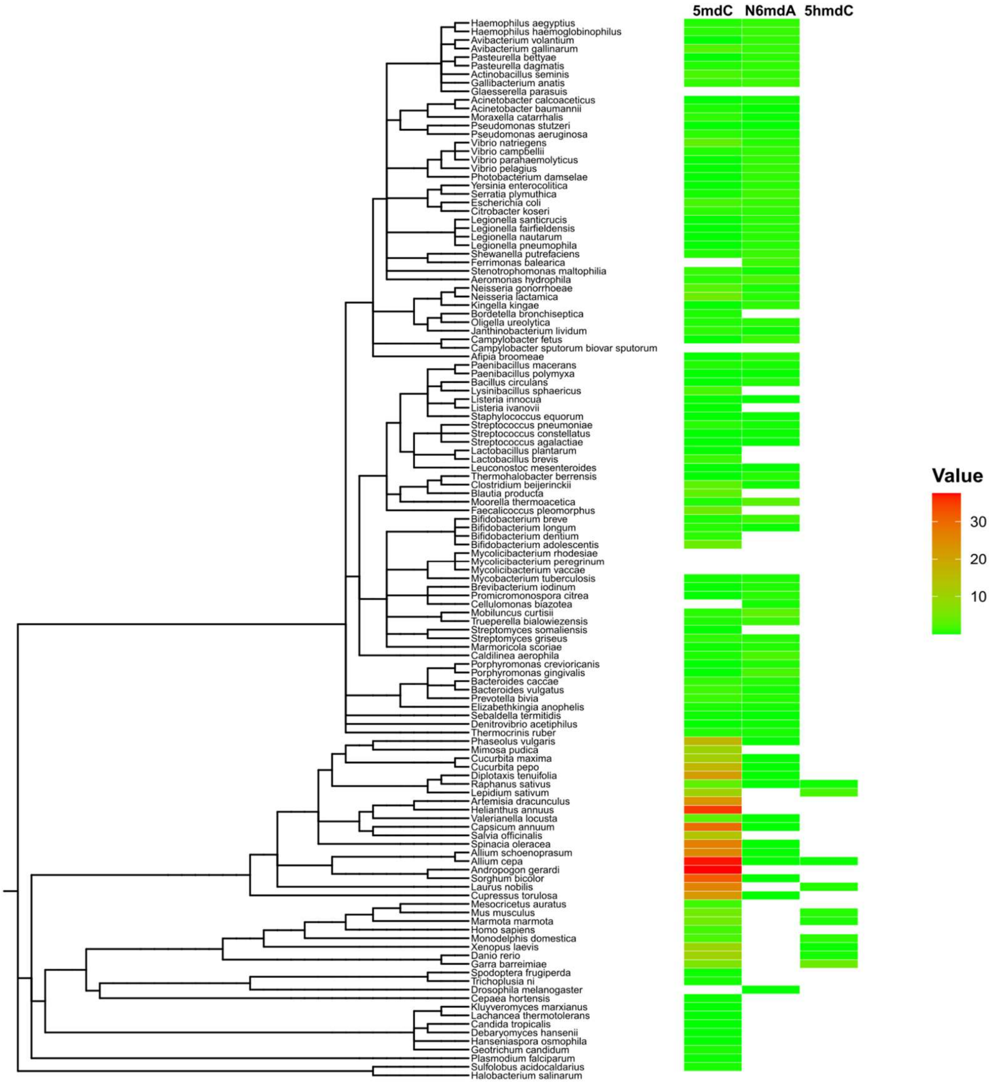


**Figure S2**: Distribution of modifications across species.

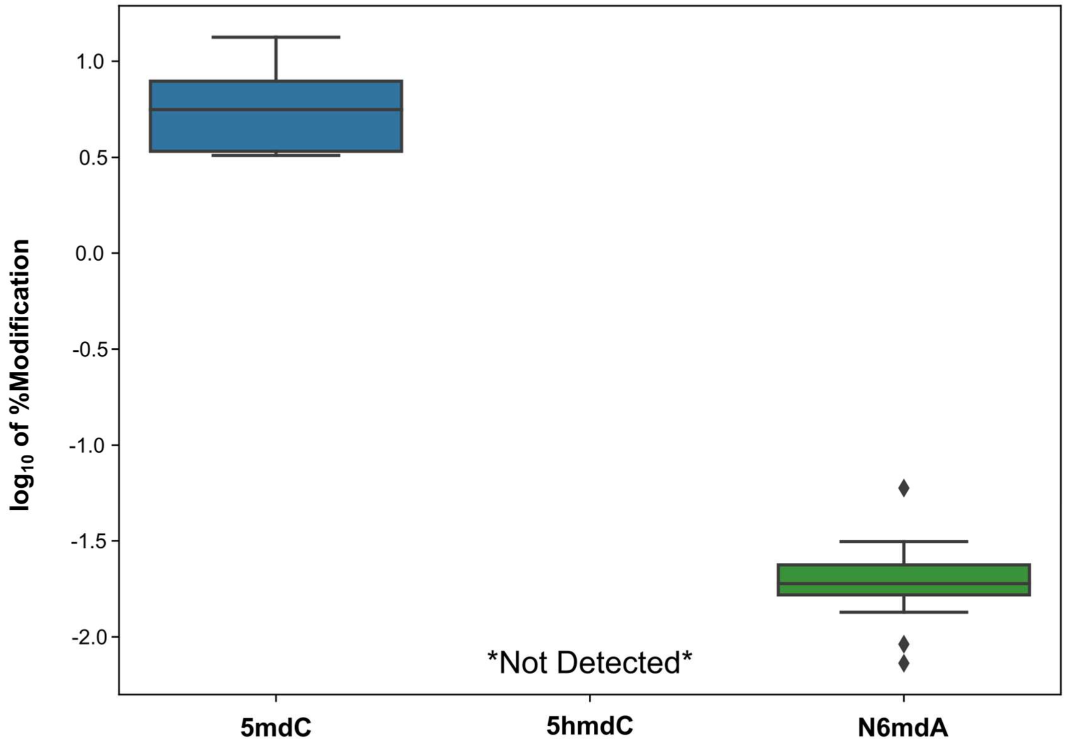


**Figure S3:** Variation in % modification (as $\log_{10}$) in HeLa cells cultured under different growth environments: (1) standard media, (2) media with no penicillin/streptomycin, (3) media with 10x penicillin/streptomycin concentration, (4) incubation at 40 °C, (5) treatment with 2.5 mM DTT, (6) treatment with 200 μM $H_2O_2$ and interferon gamma treatment. 5hmdC was below the detection limit for all samples.

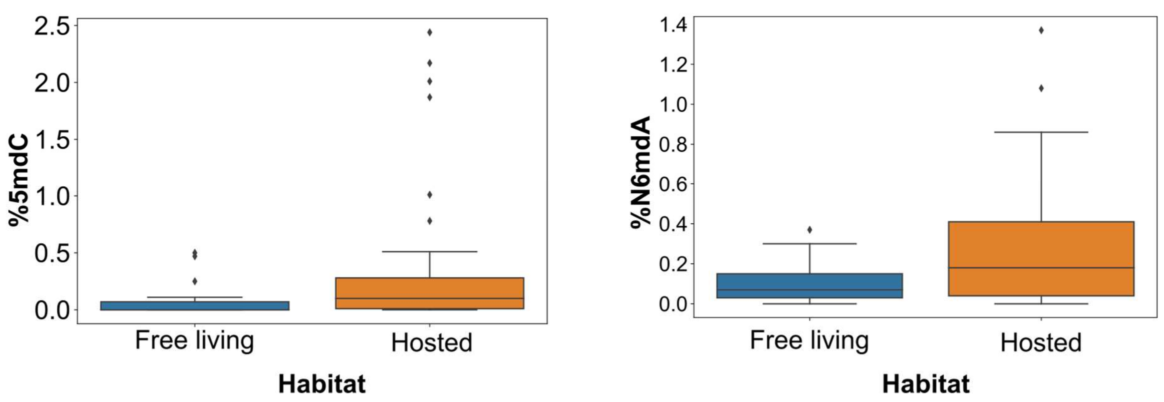

**Figure S4**: Variation of 5mdC and N6mdA among hosted and free-living bacteria.

**Table S4:** Percentages of 5mdC (wrt dG), 5hmdC (wrt dG), and N6mdA (wrt T) of different bacterial samples measured and their habitats*. Information regarding bacterial habitats was retrieved from the bacterial metadatabase BacDive (http://bacdive.dsmz.de, Accessed 14 April, 2020).(Reimer et al. 2019)

| **Name** | **%5mdC** | **%5hmdC** | **%N6mdA** | **Habitat** |
|---|---|---|---|---|
| *Acinetobacter calcoaceticus* | 0 | 0 | 0.1 | Free living |
| *Actinobacillus seminis* | 0.78 | 0 | 0.34 | Hosted |
| *Aeromonas hydrophila* | 0.13 | 0 | 0.41 | Hosted |
| *Afipia broomeae* | 0.01 | 0 | 0.1 | Hosted |
| *Avibacterium gallinarum* | 1.01 | 0 | 0.39 | Hosted |
| *Avibacterium volantium* | 0 | 0 | 0.41 | Hosted |
| *Bacillus circulans* | 0 | 0 | 0.07 | Free living |
| *Bacteroides caccae* | 0.16 | 0 | 0.09 | Hosted |
| *Bifidobacterium adolescentis* | 1.87 | 0 | 0 | Hosted |
| *Bifidobacterium breve* | 0.16 | 0 | 0.62 | Hosted |
| *Bifidobacterium dentium* | 0.26 | 0 | 0 | Hosted |
| *Bifidobacterium longum* | 0.25 | 0 | 0.04 | Hosted |
| *Blautia producta* | 0.51 | 0 | 0 | Hosted |

| | | | | |
|---|---|---|---|---|
| *Bordetella bronchiseptica* | 0.07 | 0 | 0 | Hosted |
| *Brevibacterium iodinum* | 0.02 | 0 | 0.12 | Hosted |
| *Caldilinea aerophila* | 0.07 | 0 | 0.37 | Free living |
| *Campylobacter fetus* | 0.01 | 0 | 0.62 | Hosted |
| *Campylobacter sputorum* | 0.01 | 0 | 0.86 | Hosted |
| *Cellulomonas biazotea* | 0 | 0 | 0.05 | Free living |
| *Clostridium beijerinckii* | 0.5 | 0 | 0.03 | Free living |
| *Denitrovibrio acetiphilus* | 0 | 0 | 0.04 | Free living |
| *Elizabethkingia anophelis* | 0.07 | 0 | 0.05 | Hosted |
| *Faecalicoccus pleomorphus* | 2.17 | 0 | 0 | Hosted |
| *Ferrimonas balearica* | 0 | 0 | 0.3 | Free living |
| *Gallibacterium anatis* | 0.37 | 0 | 0.52 | Hosted |
| *Haemophilus aegyptius* | 0.23 | 0 | 0.48 | Hosted |
| *Haemophilus haemoglobinophilus* | 0.28 | 0 | 0.41 | Hosted |
| *Haemophilus parasuis* | 0 | 0 | 0.42 | Hosted |
| *Kingella kingae* | 0 | 0 | 0.41 | Hosted |
| *Lactobacillus brevis* | 0.47 | 0 | 0 | Free living |

| | | | | |
|---|---|---|---|---|
| *Legionella fairfieldensis* | 0 | 0 | 0.21 | Free living |
| *Legionella nautarum* | 0 | 0 | 0.16 | Free living |
| *Legionella pneumophila* | 0 | 0 | 0.16 | Hosted |
| *Legionella santicrucis* | 0 | 0 | 0.13 | Free living |
| *Leuconostoc mesenteroides* | 0.01 | 0 | 0.03 | Free living |
| *Listeria innocua* | 0.02 | 0 | 0.01 | Free living |
| *Lysinibacillus sphaericus* | 0.38 | 0 | 0 | Hosted |
| *Marmoricola scoriae* | 0.04 | 0 | 0.13 | Free living |
| *Mobiluncus curtisii* | 0.3 | 0 | 1.37 | Hosted |
| *Moorella thermoacetica* | 0.12 | 0 | 1.08 | Hosted |
| *Moraxella catarrhalis* | 0.41 | 0 | 0.05 | Hosted |
| *Mycobacterium peregrinum* | 0.1 | 0 | 0.07 | Hosted |
| *Mycobacterium rhodesiae* | 0 | 0 | 0.01 | Hosted |
| *Mycobacterium vaccae* | 0 | 0 | 0.02 | Hosted |
| *Neisseria lactamica* | 2.01 | 0 | 0.26 | Hosted |
| *Neisseria gonorrhoeae* | 1.39 | 0 | 0.16 | Hosted |
| *Oligella ureolytica* | 0.15 | 0 | 0.25 | Hosted |

| | | | | |
|---|---|---|---|---|
| *Paenibacillus macerans* | 0.05 | 0 | 0.03 | Free living |
| *Paenibacillus polymyxa* | 0 | 0 | 0.01 | Free living |
| *Pasteurella bettyae* | 0.01 | 0 | 0.31 | Hosted |
| *Pasteurella dagmatis* | 0.21 | 0 | 0.37 | Hosted |
| *Photobacterium damselae* | 0 | 0 | 0.18 | Hosted |
| *Porphyromonas crevioricanis* | 0.01 | 0 | 0.19 | Hosted |
| *Porphyromonas gingivalis* | 0.01 | 0 | 0.81 | Hosted |
| *Prevotella bivia* | 0.49 | 0 | 0.2 | Hosted |
| *Pseudomonas stutzeri* | 0 | 0 | 0.01 | Hosted |
| *Sebaldella termitidis* | 2.44 | 0 | 0.1 | Hosted |
| *Serratia plymuthica* | 0.04 | 0 | 0.26 | Hosted |
| *Shewanella putrefaciens* | 0.08 | 0 | 0.25 | Hosted |
| *Staphylococcus equorum* | 0 | 0 | 0.03 | Free living |
| *Stenotrophomonas maltophilia* | 0.23 | 0 | 0.04 | Hosted |
| *Streptococcus agalactiae* | 0.01 | 0 | 0 | Hosted |
| *Streptomyces griseus* | 0.11 | 0 | 0.07 | Free living |
| *Streptomyces somaliensis* | 0 | 0 | 0 | Hosted |

| | | | | |
|---|---|---|---|---|
| *Thermocrinis ruber* | 0.25 | 0 | 0.15 | Free living |
| *Thermohalobacter berrensis* | 0 | 0 | 0.17 | Free living |
| *Trueperella bialowiezensis* | 0.18 | 0 | 0.56 | Hosted |
| *Vibrio campbellii* | 0.08 | 0 | 0.14 | Free living |
| *Vibrio parahaemolyticus* | 0 | 0 | 0.15 | Hosted |
| *Yersinia enterocolitica* | 0.02 | 0 | 0.13 | Hosted |

*Percentages are normalized to genome size.

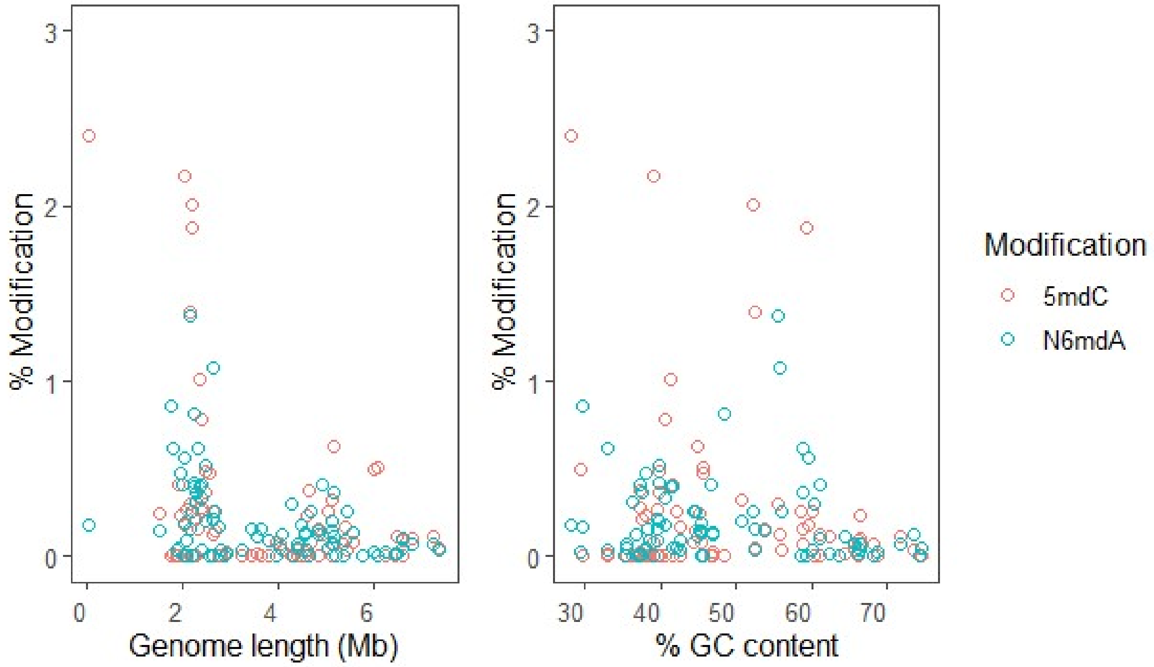


**Figure S5:** Variation of 5mdC and N6mdA according to genome length and % GC content.
